# An epigenetic predictor of death captures multi-modal measures of brain health

**DOI:** 10.1101/703504

**Authors:** Robert F. Hillary, Anna J. Stevenson, Simon R. Cox, Daniel L. McCartney, Sarah E. Harris, Anne Seeboth, Jon Higham, Duncan Sproul, Adele M. Taylor, Paul Redmond, Janie Corley, Alison Pattie, Maria del. C Valdés Hernández, Susana Muñoz-Maniega, Mark E. Bastin, Joanna M. Wardlaw, Steve Horvath, Craig W. Ritchie, Tara L. Spires-Jones, Andrew M. McIntosh, Kathryn L. Evans, Ian J. Deary, Riccardo E. Marioni

## Abstract

Individuals of the same chronological age exhibit disparate rates of biological ageing. Consequently, a number of methodologies have been proposed to determine biological age and primarily exploit variation at the level of DNA methylation (DNAm) – a commonly studied epigenetic mechanism. A novel epigenetic clock, termed ‘DNAm GrimAge’ has outperformed its predecessors in predicting the risk of mortality as well as a number of age-related morbidities. However, the association between DNAm GrimAge and cognitive or neuroimaging phenotypes remains unknown. We explore these associations in the Lothian Birth Cohort 1936 (n=709, mean age 73 years). Higher DNAm GrimAge was strongly associated with all-cause mortality over twelve years of follow-up (Hazard Ratio per standard deviation increase in GrimAge: 1.81, P < 2.0 × 10^-16^). Higher DNAm GrimAge was associated with lower age 11 IQ (β=-0.11), lower age 73 general cognitive ability (β=-0.18), decreased brain volume (β=-0.25) and increased brain white matter hyperintensities (β=0.17). Sixty-eight of 137 health- and brain-related phenotypes tested were significantly associated with DNAm GrimAge. Adjusting all models for childhood cognitive ability attenuated to non-significance a small number of associations (12/68 associations; 6 of which were cognitive traits), but not the association with general cognitive ability (33.9% attenuation). Higher DNAm GrimAge cross-sectionally associates with lower cognitive ability and brain vascular lesions in older age, independently of early life cognitive ability. Thus, this epigenetic predictor of mortality is also associated with multiple different measures of brain health and may aid in the prediction of age-related cognitive decline.

## 1. Introduction

The rapid ageing of the global population has resulted in an increase in the personal and societal burden of age-associated disease and disability [1]. Consequently, there is an urgent need to identify those individuals at high risk of age-related morbidities and mortality to allow for early intervention. Recently, a number of methods for determining biological age have been developed which leverage inter-individual variation in physiological and molecular characteristics [2-6]. Primarily, these measures of biological age have focussed on variation at the level of DNA methylation (DNAm). DNAm is a commonly-studied epigenetic mechanism typically characterised by the addition of a methyl group to a cytosine-phosphate-guanine (CpG) nucleotide base pairing, thereby permitting regulation of gene activity [7]. Crucially, these biological age predictors, also referred to as ‘epigenetic clocks’, correlate strongly with chronological age; furthermore, for a given chronological age, an advanced epigenetic age is associated with increased mortality risk and many age-related morbidities [8-12].

A novel epigenetic clock, termed ‘DNAm GrimAge’ has been developed to predict mortality [13]. To derive DNAm GrimAge, an elastic net Cox regression model was used to regress time-to-death due to all-cause mortality on chronological age, sex and DNAm-based surrogates for smoking pack years and 12 plasma proteins. The model selected chronological age, sex, and methylation-based surrogates for smoking pack years and for 7/12 plasma proteins. The linear combination of these variables allows for an estimation of DNAm GrimAge. As with other epigenetic clocks, if an individual’s DNAm GrimAge is higher than their chronological age, then this provides a measure of accelerated biological ageing. Lu *et al.* (2019) comprehensively demonstrated that an accelerated DNAm GrimAge (also known as AgeAccelGrim) is associated with a number of peripheral, lifestyle and cardiometabolic traits and outperforms predecessor clocks in predicting death. However, the relationship between an accelerated GrimAge and cognitive, and neuroimaging, phenotypes remains unexplored. As brain structure and cognitive function show mean declines with age, and associate with disability and disease burden, the discovery of molecular correlates of neurological and neurostructural aberrations may be of particular benefit in gerontology [14, 15]. Therefore, in this study, we test the hypothesis that, in a large, narrow age-range population cohort of older adults (Lothian Birth Cohort 1936 (LBC1936)), an accelerated DNAm GrimAge is cross-sectionally associated with poorer cognitive performance, structural neuroimaging measures, and neurology-related proteins.

Additionally, higher childhood intelligence (as defined by age 11 IQ) is associated with a lower risk of mortality across the life course [16-18]. Furthermore, childhood intelligence associates with a healthier lifestyle and less morbidity in middle age, as well as a lower allostatic load in older age [19-21]. Furthermore, intelligence in early life is related to variability in cortical thickness, white matter macro- and micro-structure, as well as cognitive ability, fewer vascular lesions and lower risk of stroke in later life [22-27]. Notably, adjustment for age 11 IQ was recently shown to attenuate associations between another epigenetic clock measure, DNAm PhenoAge, with a wide range of phenotypes including cognitive traits in LBC1936 [28]. Therefore, we also test the hypothesis that controlling for childhood intelligence attenuates associations between DNAm GrimAge and mortality, cognitive and neuroimaging measures, as well as neurology-related proteins in older age.

## 2. Materials and Methods

### 2.1 The Lothian Birth Cohort 1936

The LBC1936 comprises Scottish individuals born in 1936, most of whom took part in the Scottish Mental Survey 1947 at age 11. Participants who were living within Edinburgh and the Lothians were re-contacted approximately 60 years later. Of these participants, 1,091 consented and joined the LBC1936. Upon recruitment, participants were approximately 70 years of age (mean age: 69.6 ± 0.8 years) and subsequently attended four additional waves of clinical examinations about every three years. Detailed genetic, epigenetic, physical, psychosocial, cognitive, neuroimaging, health and lifestyle data are available for members of the LBC1936. Recruitment and testing of the LBC1936 have been described previously [29, 30].

### 2.2 Methylation preparation in the Lothian Birth Cohort 1936

DNA from whole blood was assessed using the Illumina 450K methylation array at the Edinburgh Clinical Research Facility. Details of quality control procedures have been described elsewhere (see Supplementary Methods) [31, 32].

### 2.3 Derivation of DNAm GrimAge

DNAm GrimAge was calculated using the online age calculator (https://dnamage.genetics.ucla.edu/) developed by Horvath [33]. LBC1936 methylation data were used as input for the algorithm and data underwent a further round of normalisation by the age calculator. The DNAm GrimAge biomarker was calculated using a method developed by Lu *et al* [13] and is based on a linear combination of age, sex, DNAm-based surrogates for smoking, and seven proteins (adrenomedulin (DNAm ADM), beta-2-microglobulin (DNAm B2M), cystatin C (DNAm Cystatin C), growth differentiation factor 15 (DNAM GDF-15), leptin (DNAm leptin), plasminogen activation inhibitor 1 (DNAm PAI-1), and tissue inhibitor metalloproteinaise (DNAm TIMP-1)). Supplementary Figure 1 shows the correlation between all methylation-based surrogates. All predictors, with the exception of DNAm Leptin (r^2^ = - 0.29), were positively correlated with DNAm GrimAge (absolute range = [0.24: 0.82], median = 0.25 and mean of correlation coefficients = 0.25). The difference between DNAm GrimAge and chronological age (an accelerated DNAm GrimAge) provides a measure of biological ageing. In a previous study, for a given chronological age, individuals with higher DNAm GrimAge had a higher risk for mortality than individuals of the same chronological age with a lower DNAm GrimAge [13].

**Figure 1.**
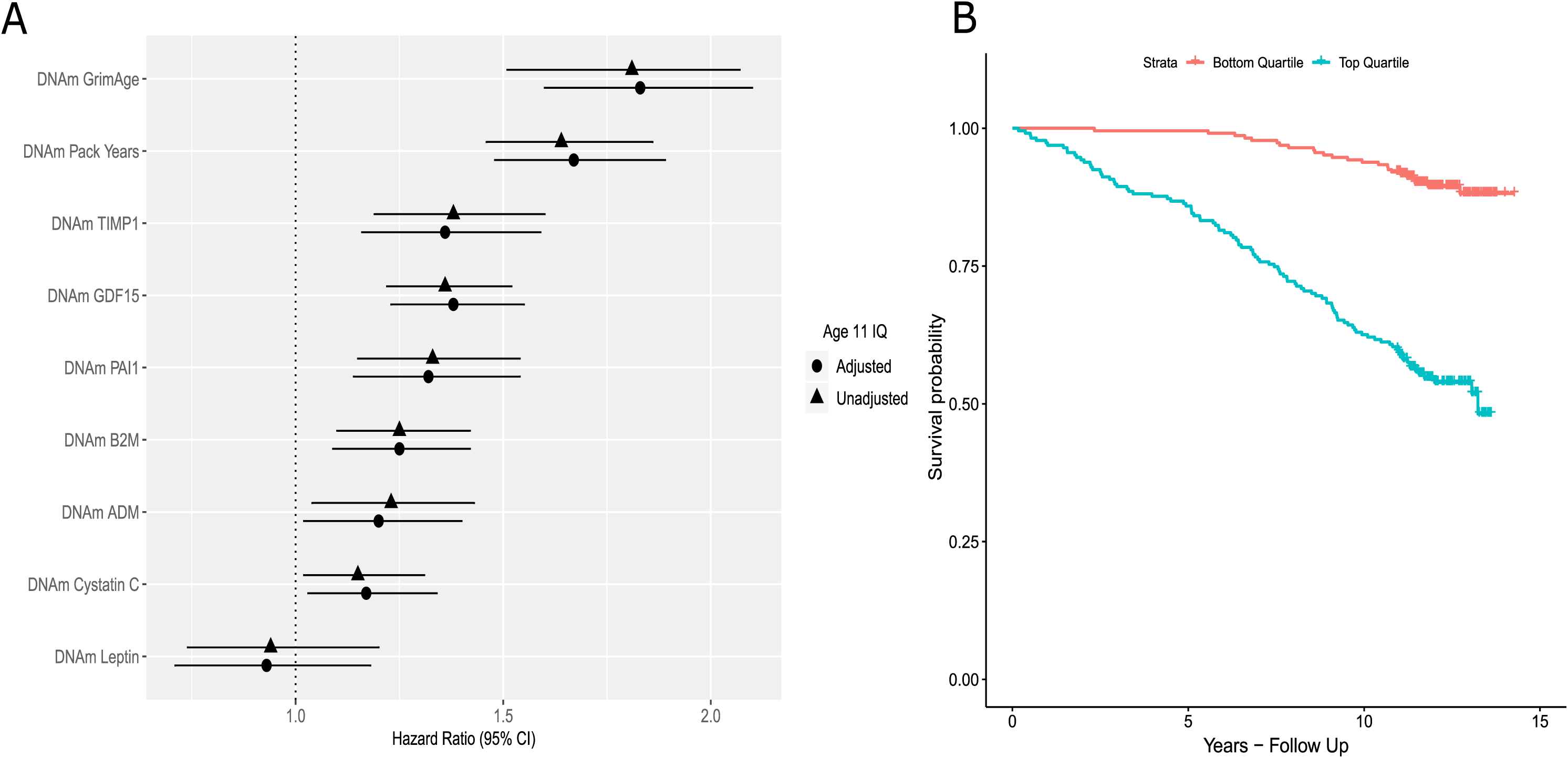
DNAm GrimAge and its component surrogate markers predict mortality in the LBC1936. (A) Forest plot showing hazard ratios and 95% confidence intervals (horizontal lines) from Cox proportional hazard models for DNAm GrimAge and its constituent DNAm surrogate markers in the LBC1936 (n = 906, no. of deaths = 226 following 12 years of follow-up). All associations with the exceptions of DNAm Leptin were significant. (B) Kaplan-Meier survival curve exhibiting the survival probabilities for the top (highest DNAm GrimAge) and bottom quartiles (lowest DNAm GrimAge) for DNAm GrimAge in the LBC1936 following 12 years of-follow up.

### 2.4 Phenotypic Data

Our phenotypic analyses were divided into four sections. Firstly, we examined the association between age-adjusted DNAm GrimAge and mortality in the LBC1936 over 12 years of follow-up. For our survival models, we used measures of age-adjusted DNAm GrimAge at Wave 1 of the LBC1936 study (n = 906 individuals; age: 70 years). For all other phenotypic analyses, we examined cross-sectional associations with age-adjusted DNAm GrimAge at Wave 2 (age: 73 years) as complete proteomic, DNA methylation and phenotypic data were available at this time point (n = 709 individuals). We also investigated the cross-sectional association of an accelerated DNAm GrimAge with a number of physical (body mass index, height, grip strength, lung function and weight) and blood traits (albumin, C-reactive protein, cholesterol, creatinine, ferritin, interleukin-6 and iron; at Wave 2; age 73 years) that have been related to mortality and frailty in older age [34-42].

Secondly, we tested the association between an accelerated DNAm GrimAge and cognitive traits (n = 18 phenotypes). Cognitive tests taken at Wave 2 (age: 73 years) included six Wechsler Adult Intelligence Scale-III UK (WAIS-III) non-verbal subtests (matrix reasoning, letter number sequencing, block design, symbol search, digit symbol, and digit span backward). Principal component analysis (PCA) was performed using these cognitive tests and scores on the first un-rotated principal component (general cognitive ability, *g*) were extracted which explained 51% of variance. Individual test loadings ranged from 0.65 to 0.75. Wechsler Memory Scale-III items as well as measures of crystallised intelligence and reaction time were also examined in relation to DNAm GrimAge. Additionally, we examined whether an accelerated DNAm GrimAge associated with *APOE* ε4 carrier status.

Thirdly, we tested the association between an accelerated DNAm GrimAge and neuroimaging phenotypes at Wave 2 (age: 73 years, see Supplementary Methods). The brain MRI acquisition and processing pipeline has been made available in an open access protocol paper [43]. Total brain, normal-appearing white matter, grey matter and white matter hyperintensity volumes were segmented using a semi-automated multi-spectral technique [44]. These volumes were then expressed as a proportion of intracranial volume (ICV), which controls for the confounding effect of head size. The resultant ratios were tested for associations with age-adjusted DNAm GrimAge. Diffusion-tensor imaging-derived measures of fractional anisotropy (FA) and mean diffusivity (MD) were obtained for participants at Wave 2 (age: 73 years). Prior to conducting region-specific analyses, general factors of FA (gFA) and MD (gMD) were derived by entering the left and right FA and MD values of each tract separately into a PCA. Scores from the first un-rotated principal component were extracted and labelled as gFA (variance explained: 52%, loadings: 0.46 – 0.95) or gMD (variance explained: 48%, loadings: 0.47 – 0.88), respectively. These general factors reflect common microstructural properties across main white matter pathways and capture the common variance in white matter integrity [45].

Fourthly, we tested the association between an accelerated DNAm GrimAge and the levels of 92 neurological protein biomarkers (Olink^®^ neurology panel). The neurology panel represents proteins with established links to neuropathology as well as exploratory proteins with roles in processes including cellular communication and immunology. Plasma was extracted from 816 blood samples collected in citrate tubes at mean age 72.5 ± 0.7 years (Wave 2; Supplementary Methods). Protein levels were transformed by rank-based inverse normalisation. Normalised protein levels were regressed onto age-adjusted DNAm GrimAge.

Descriptive statistics for phenotypes are presented in Supplementary File 1. Data collection protocols have been described fully previously and are described in Supplementary Note 1 [46].

### 2.5 Statistical analyses

DNAm GrimAge was regressed onto chronological age for all LBC1936 participants. These residuals were defined as an accelerated DNAm GrimAge (also known as AgeAccelGrim). Linear regression models were used to investigate relationships between continuous variables and an accelerated DNAm GrimAge, as well as age-adjusted methylation-based surrogates for smoking pack years and the plasma proteins that feed into DNAm GrimAge. Logistic regression was used to test the association between methylation-based predictors and *APOE* ε4 carrier status. An accelerated DNAm GrimAge, age-adjusted DNAm Pack Years or age-adjusted DNAm plasma protein levels were the independent variable of interest in each regression model and all variables were scaled to have a mean of zero and unit variance. Height and smoking status were included as covariates in the models for lung function (forced expiratory volume FEV1; forced vital capacity: FVC; forced expiratory ratio: FER; and peak expiratory flow: PEF). All models were adjusted for chronological age and sex. To investigate possible statistical confounding by childhood cognitive ability, all models were repeated with adjustment for age 11 IQ scores. To correct for multiple testing, and given that the methylation-based predictors exhibited a high degree of inter-correlation, we applied the false discovery rate (FDR; [47]) method to phenotypic association analyses (n = 137 phenotypes), separately for each predictor. Associations between age-adjusted DNAm GrimAge and regional cortical volume were conducted using the SurfStat toolbox (http://www.math.mcgill.ca/keith/surfstat) for Matrix Laboratory R2018a (The MathWorks Inc, Natick, MA), using the same covariates as above and FDR correction for multiple testing.

### 2.6 Ethics and consent

Ethical permission for LBC1936 was obtained from the Multi-Centre Research Ethics Committee for Scotland (MREC/01/0/56), the Lothian Research Ethics Committee (Wave 1: LREC/2003/2/29) and the Scotland A Research Ethics Committee (Waves 2, 3 and 4: 07/MRE00/58). Written informed consent was obtained from all participants.

## 3. Results

### 3.1 Cohort characteristics

Details of LBC1936 participant characteristics at Waves 1 and 2 are presented in Supplementary File 1. Briefly, 47.6% of participants in this study were female. At Wave 1 (relating to the mortality analysis), mean chronological age for both males and females was 69.6 years (SD 0.8) whereas the mean DNAm GrimAge was 67.4 years (SD 5.2). At Wave 2 (relating to cross-sectional analyses), mean chronological age for both males and females was 72.5 years (SD 0.7) whereas the mean DNAm GrimAge was 70.0 years (SD 4.9). The lower mean measure of epigenetic age when compared to chronological age may reflect overall good health of the cohort. However, the variance associated with DNAm GrimAge is much higher than that of chronological age. Mean age 11 IQ scores were 100.69 (SD: 15.37). Notably, lower IQ scores at age 11 (β = -0.11, P = 0.02) were associated with an accelerated DNAm GrimAge. Associations between age 11 IQ and tested phenotypes are presented in Supplementary File 2.

### 3.2 DNAm GrimAge predicts mortality and associates with frailty factors in the LBC1936

Mortality in LBC1936 participants was assessed in relation to an accelerated DNAm GrimAge as well as age-adjusted DNAm-based surrogate markers for plasma protein levels and smoking pack years. DNAm GrimAge was derived for 906 participants with methylation data (at Wave 1: age 70 years). There were 226 deaths (24.9%) over 12 years of follow-up.

A higher DNAm GrimAge was significantly associated with risk of all-cause mortality (Hazard Ratio (HR) = 1.81 per SD increase in DNAm GrimAge, 95% confidence interval (CI) = [1.58, 2.07], P < 2.0 × 10^-16^). Furthermore, higher levels of age-adjusted DNAm Pack Years were associated with all-cause mortality in the LBC1936 (HR = 1.64 per SD, 95% CI [1.46, 1.86], P = 2.0 × 10^-16^). In relation to methylation-based surrogates for plasma protein levels, six of the seven DNAm protein surrogates (DNAm ADM, B2M, Cystatin C, GDF15, PAI1 and TIMP1) were significantly associated with all-cause mortality (see Supplementary File 3; Figure 1A). Following adjustment for age 11 IQ, there was very little change in the HRs and all of the predictors remained significant. Indeed, hazard ratios from all survival models ranged from an attenuation of 2.4% to an increase of 1.8% following adjustment for childhood intelligence.

A Kaplan-Meier survival plot for an accelerated DNAm GrimAge, split into the highest and lowest quartiles, is presented in Figure 1B illustrating the higher mortality risk for those with a higher DNAm GrimAge. Kaplan-Meier survival plots for methylation-based surrogates for smoking pack years and plasma protein levels are presented in Supplementary Figure 2.

**Figure 2.**
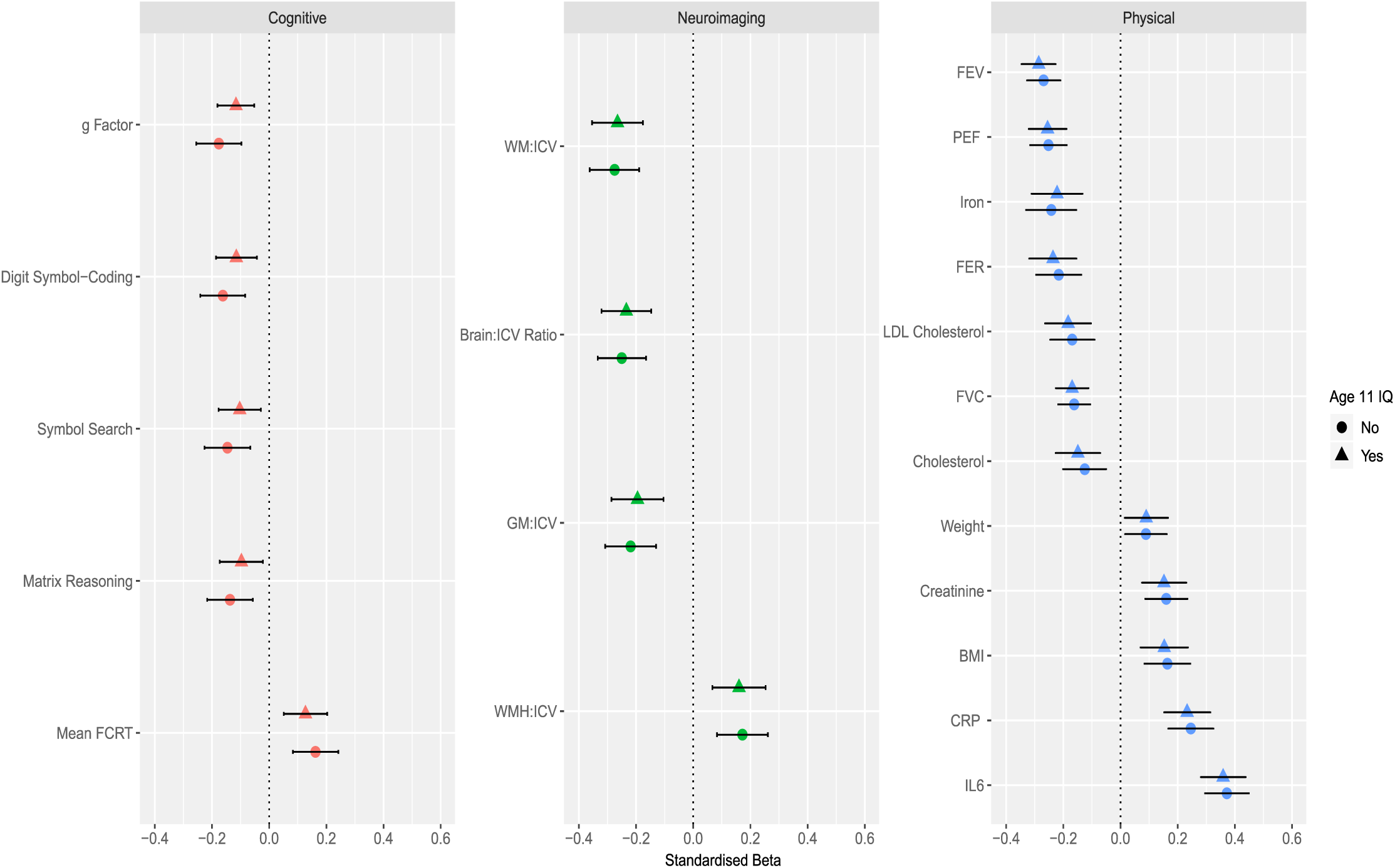
Cross-sectional association between age-adjusted DNAm GrimAge and cognitive, neuroimaging and physical traits in the LBC1936. *Cognitive*: An accelerated DNAm GrimAge was negatively associated with the general factor of cognitive ability, digit symbol-coding, symbol search and matrix reasoning tasks. DNAm GrimAge was also associated with an increased mean four choice reaction time. *Neuroimaging*: Age-adjusted DNAm GrimAge was negatively associated with the ratios of white matter volume, brain volume and grey matter volume to intracranial volume, and positively associated with volume of white matter hyperintensities to intracranial volume. *Physical:* An accelerated DNAm GrimAge was negatively associated with four measures of lung function: forced expiratory volume in 1 second, forced vital capacity, forced expiratory ratio and peak expiratory flow, as well as levels of iron, low-density lipoprotein cholesterol and total cholesterol. Age-adjusted DNAm GrimAge was positively associated with weight, levels of creatinine, body mass index as well as levels of C-reactive protein and interleukin-6. BMI (body mass index), CRP (C-reactive protein), FCRT (four choice reaction time), FER (forced expiratory ratio), FEV (forced expiratory volume), FVC (forced vital capacity), GM (grey matter), ICV (intracranial volume), IL6 (interleukin-6), LDL (low-density lipoprotein), PEF (peak expiratory flow), WM (white matter), WHM (white matter hyperintensities).

For the remainder of the results, only those associations with an FDR-corrected significant P value (<0.05) are presented herein and in Figure 2. Full results are presented in Supplementary File 4. In relation to major mortality- and frailty-associated physical traits in the LBC1936, an accelerated DNAm GrimAge was associated with increased levels of interleukin-6 (β = 0.37, P = 2.3 × 10^-18^), C-reactive protein (β = 0.25, P = 2.8 × 10^-8^), creatinine (β = 0.16, P = 1.1 × 10^-4^), an increased body mass index (β = 0.16, P = 2.9 × 10^-4^), triglyceride concentration (β = 0.13, P = 5.0 × 10^-3^) and body weight (β = 0.09, P = 0.04) (Figure 2). The relationship between accelerated DNAm GrimAge and triglycerides was no longer significant after controlling for childhood cognitive ability with the effect size decreasing from 0.13 to 0.09 (32.5% attenuation) (Supplementary File 4).

An accelerated DNAm GrimAge was negatively associated with all four measures of lung function (β = [-0.16 to -0.27], P = [9.4× 10^-7^ to 1.7 × 10^-16^]), iron levels (β = -0.24, P = 7.2 × 10^-7^), low-density lipoprotein cholesterol levels (β = -0.17, P = 1.1 × 10^-4^), total cholesterol levels (β = -0.13, P = 1.1 × 10^-4^) and height (β = -0.08, P = 0.01) (Figure 2). Only the relationship between accelerated DNAm GrimAge and height was non-significant after controlling for childhood intelligence, with the effect size attenuating from -0.08 to -0.06 (% attenuation: 24.5%) (Supplementary File 4). On average, associations were attenuated by 2.5% after controlling for age 11 IQ [ranged from: 19.1% increase (total cholesterol) to 32.5% attenuation (triglycerides)]. Relationships between all phenotypes tested in this study and age-adjusted DNAm Pack Years as well as age-adjusted plasma protein levels are presented in Supplementary File 5.

### 3.3 DNAm GrimAge associates with lower cognitive ability in the LBC1936

An accelerated DNAm GrimAge was significantly associated with lower measures of general cognitive ability (g: β = -0.18, P = 8.0 × 10^-6^; n = 709). Furthermore, an accelerated DNAm GrimAge was negatively associated with all six component tests for fluid intelligence from which g was derived (see Methods 2.4; β = [-0.11 to -0.16], P = [0.02 to 2.4 × 10^-4^]). Additionally, an accelerated DNAm GrimAge was associated with an increased four choice reaction time mean (β = 0.16, P = 2.9 × 10^-4^). Lower IQ scores at age 70 (which correlated 0.70 with age 11 IQ scores) were associated with age-adjusted DNAm GrimAge (β = -0.11, P = 0.02). An accelerated DNAm GrimAge was also negatively associated with the following measures of crystallised intelligence: the Wechsler Test of Adult Reading (β = -0.13, P = 4.0 × 10^-3^) and the National Adult Reading Test (β = -0.10, P = 0.03).

Following adjustment for age 11 IQ, an accelerated DNAm GrimAge remained significantly associated with general cognitive ability (g: β = -0.12, P = 2.0 × 10^-3^; 33.9% attenuation). Three out of the six tests which constitute the general intelligence factor remained significant after adjustment for age 11 IQ (digit-symbol coding, symbol search, and matrix reasoning). Furthermore, the association between an accelerated DNAm GrimAge and an increased mean four choice reaction time remained significant following adjustment for age 11 IQ (Figure 2). On average, associations between cognitive tasks and an accelerated DNAm GrimAge were attenuated by 41.1% following controlling for age 11 IQ (ranging from 21.7% attenuation [four choice reaction time] to 77.4% attenuation [National Adult Reading Test]). All associations between cognitive traits and an accelerated DNAm GrimAge in this study are presented in Supplementary Figure 4. Finally, an accelerated DNAm GrimAge was not associated with *APOE* ε4 carrier status – the strongest genetic risk factor for Alzheimer’s disease (Odds Ratio = 0.96, 95% CI = [0.93, 1.00], P = 0.06).

### 3.4 DNAm GrimAge is associated with gross neurostructural differences in the LBC1936

An accelerated DNAm GrimAge was associated with lower white matter volume (β = -0.28, P = 1.7 × 10^-8^), total brain volume (β = -0.25, P = 1.4 × 10^-7^) and grey matter volume (β = -0.22, P = 1.3 × 10^-5^). Furthermore, an accelerated DNAm GrimAge was associated with an increased volume of white matter hyperintensities (β = 0.17, P = 1.0 × 10^-3^) (Figure 2). All associations remained significant following adjustment for age 11 IQ (Supplementary File 4). On average, these associations were attenuated by 6.98% after adjusting for age 11 IQ. All associations between neuroimaging traits and an accelerated DNAm GrimAge in this study are presented in Supplementary Figure 5.

An accelerated DNAm GrimAge was not significantly associated with general factors of white matter microstructural metrics i.e. Fractional Anisotropy (β = -0.009, P = 0.89) or Mean Diffusivity (β = -0.001, P = 0.98), hence additional regional analyses were not performed. However, given that DNAm GrimAge was associated with grey matter volume, we further tested whether there was regional cortical heterogeneity in relation to the DNAm GrimAge-grey matter association. The negative association between accelerated DNAm GrimAge and cortical volume showed a degree of regional heterogeneity across the cortical surface (Figure 3). The strongest magnitudes were evident in frontal (superior lateral and medial) and temporal (lateral and medial) regions, motor and somatosensory cortex, and posterior cingulate/precuneal areas. In contrast, associations between DNAm GrimAge and cortical volume were predominantly non-significant in parietal, occipital, inferior lateral and ventromedial frontal regions. When the associations were additionally corrected for age 11 IQ, the magnitude of the effect sizes at the FDR-significant loci were weakly attenuated (mean t-value attenuation = 3.36%; Supplementary Figure 6).

**Figure 3.**
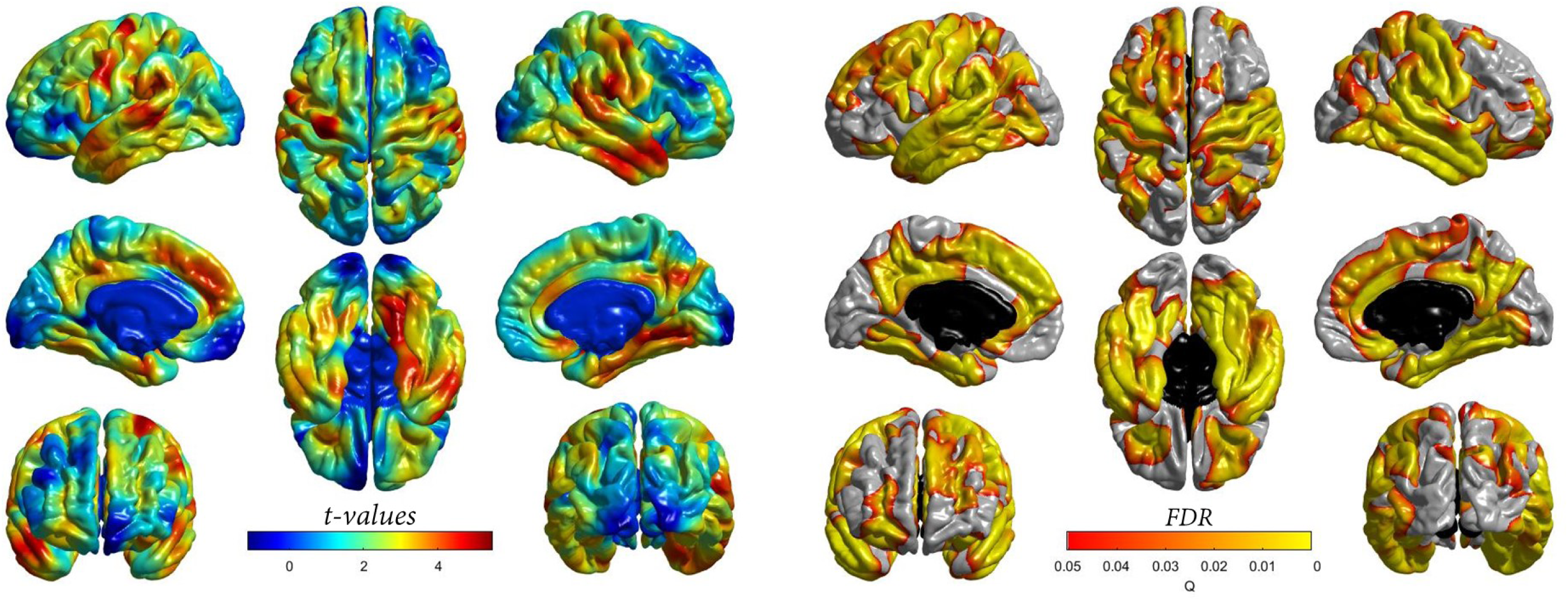
Cross-sectional association between age-adjusted DNAm GrimAge and regional cortical volume in the LBC1936. *Left panel:* t-values indicate the magnitude of the negative association (values have been flipped for visualisation purposes). An accelerated DNAm GrimAge was negatively associated with cortical volume. *Right Panel*: Corresponding FDR-corrected P values indicate the spatial distribution of significant associations. FDR (false discovery rate).

### 3.5 Association of DNAm GrimAge with neurological protein biomarkers

Forty of the 92 neurology-related Olink^®^ proteins were significantly associated with an accelerated DNAm GrimAge at FDR-corrected P < 0.05 (n = 709). These proteins explained between 0.73% (β = -0.09, NC-Dase) to 7.19% (β = 0.30, SKR3) of inter-individual variation in an accelerated DNAm GrimAge (in a model which was not adjusted for age and sex; Supplementary File 6). Following adjustment for age 11 IQ, 36/40 associations (90%) remained significant. After adjusting for age 11 IQ, associations were, on average, attenuated by 3.03%.

### 3.6 Correlation between DNAm GrimAge and DNAm Pack Years

We observed that DNAm GrimAge and DNAm Pack Years were highly correlated (correlation coefficient: 0.82) and were cross-sectionally associated with many of the same variables in our phenotypic analyses (Supplementary File 7). Therefore, we carried out a follow-up analysis to determine the difference in magnitude between the effect sizes for DNAm GrimAge or DNAm Pack Years in relation to phenotypes associated with both predictors. Prior to adjusting for age 11 IQ, the effect sizes had a correlation coefficient of 0.88. However, they were, on average, 16.5% greater for DNAm GrimAge when compared to DNAm Pack Years.

Following adjustment for age 11 IQ, the correlation coefficient was 0.84, and the effect sizes were, on average, 23.1% greater for DNAm GrimAge upon comparison to DNAm Pack Years. A plot demonstrating the correlation between effect sizes for DNAm GrimAge and DNAm Pack Years from our cross-sectional phenotypic analyses is presented in Supplementary Figure 7.

## Discussion

In this study, we found that a higher-than-expected DNAm GrimAge strongly predicted mortality and was associated with a number of mortality- and frailty-associated traits. This provides the first external replication of the association between DNAm GrimAge and survival. After controlling for childhood cognitive ability, we found that an accelerated DNAm GrimAge was cross-sectionally associated with lower general cognitive ability as well as lower scores on processing speed and perceptual organisation tasks, and slower reaction time speed; this effectively means that accelerated DNAm GrimAge was associated with more cognitive decline. Furthermore, an accelerated DNAm GrimAge was associated with gross neuroanatomical differences and vascular lesions in older age. Finally, a number of neurology-related proteins were associated with an accelerated DNAm GrimAge.

DNAm GrimAge was developed using mortality as a reference and consequently supplants its predecessors in relation to mortality risk prediction. Indeed, in this study, we observed a hazard ratio of 1.81 per standard deviation increase in an accelerated DNAm GrimAge, which outperforms that of previous epigenetic clocks (Hannum Age HR: 1.22, Horvath Age HR: 1.19; DNAm PhenoAge HR: 1.17; all applied to LBC1936) [8, 28]. In relation to mortality- and frailty-associated traits, the strongest association was between DNAm GrimAge and interleukin-6. Furthermore, DNAm GrimAge was strongly associated with C-reactive protein (whose production is stimulated by interleukin-6). Together, this corroborates evidence for the “inflammaging” theory which postulates that chronic, low-grade inflammation significantly influences biological ageing and decline [48]. An accelerated DNAm GrimAge was also associated with lower low-density lipoprotein cholesterol and total cholesterol. In older age, lower levels of these blood-based factors are also associated with higher risk of mortality [49]. Additionally, DNAm GrimAge was associated with a higher body mass index which does not agree with previous findings showing that an increased body mass index is protective against mortality risk [39]. However, this may be driven by a strong association between DNAm Leptin and body mass index. Indeed, leptin is an adipose tissue-derived hormone which acts an appetite suppressant, and is strongly correlated with body mass index and obesity [50, 51].

We observed a significant relationship between higher childhood intelligence (as well as age 70 IQ) and a lower DNAm GrimAge in older age. After controlling for childhood cognitive ability, associations between DNAm GrimAge and tests of crystallised intelligence were attenuated to non-significance. This finding is not surprising given that crystallised intelligence remains stable throughout adulthood [52], and that the National Adult Reading Test strongly retrodicts childhood IQ in this sample [53]. However, relationships between DNAm GrimAge and general cognitive ability, as well as fluid intelligence measures, remained significant after adjusting for age 11 IQ. Nevertheless, these associations were attenuated by an average of 41.4% following adjustment for age 11 IQ. Therefore, blood-based methylation changes, as captured by DNAm GrimAge, helps to explain additional variance in late life cognitive ability and fluid intelligence.

An accelerated DNAm GrimAge was significantly associated with gross neurostructural differences, including reductions in total brain, grey matter and white matter volumes and increases in white matter hyperintensity volumes. There was also some heterogeneity in the associations with regional cortical volume, whereby effects were strongest in frontal (superior lateral and medial) and temporal regions. These regions also exhibit the largest annual decrease in middle and older age [54], and are most informative for predicting chronological age (albeit using cortical thickness rather than volume; [55]). White matter hyperintensities, which associate with DNAm GrimAge, have also been linked to cortical loss in temporal and lateral frontal regions [56]. This may indicate that altered methylation profiles could help explain mechanistic relationships between neurovascular lesions and cortical atrophy. However, adjustment for vascular risk factors such as hypercholesterolemia, smoking and diabetes is merited in this context. Furthermore, white matter hyperintensities are also related to physical disability, processing speed and cognitive decline [57, 58]. Additionally, the presence of white matter hyperintensities doubles the risk of dementia, and triples the risk of stroke, and is associated with clinical outcomes in stroke [59, 60]. Therefore, DNAm GrimAge may capture vital aspects of age-related alterations in neurostructural integrity and gross brain pathology.

We observed a very strong correlation between DNAm GrimAge and DNAm Pack Years. Indeed, the associations between smoking and mortality, cognitive decline and brain pathology are well-documented [61-63]. However, the larger effect sizes for DNAm GrimAge suggest that this composite biomarker is supplemented by the inclusion of methylation-based surrogates for plasma protein levels. Additionally, we identified associations with a number of neurology-related proteins (n = 40 before adjustment for age 11 IQ; n = 36 after adjustment for age 11 IQ) which may further inform the risk of mortality and age-related morbidities, particularly in relation to neurological disease. Future studies are necessary to further define the biological relationships between such proteins and their relevance to age-related pathologies and cognitive decline.

The use of methylation-based proxies for smoking pack years and proteomic data is advantageous as methylation-based predictors are often more accurate than self-reported phenotypes, and the cost of complex proteomic platforms is negated [64]. One strength of this study is that rich data were available across the eighth decade of life, a period in which risk of cognitive decline and compromised brain integrity increases significantly. However, LBC1936 comprises relatively healthy older adults, complicating the generalisability of findings to at-risk clinical populations and broader age ranges.

In conclusion, we demonstrated that an epigenetic predictor of mortality associates with cognitive ability, cognitive decline and neuroimaging phenotypes in a cohort of healthy older ageing adults. These associations were largely independent of another well-known predictor of mortality, childhood intelligence. Indeed, methylation alterations in blood, as captured by DNAm GrimAge, could help provide early indications towards mortality prediction and decline in brain health.

## Acknowledgements

We thank the LBC1936 participants and study team. We thank the nursing staff at the Wellcome Trust Clinical Research Facility, and radiographers at the Brain Research Imaging Research Centre (www.bric.ed.ac.uk), Western General Edinburgh. The LBC1936 is funded by Age UK as The Disconnected Mind project, and by the Medical Research Council (G0701120, G1001245, MR/M013111/1, MR/R024065/1). S.R.C, M.E.B and I.J.D are also supported by a National Institutes of Health (NIH) research grant R01AG054628. R.F.H and A.J.S are supported by funding from the Wellcome Trust 4-year PhD in Translational Neuroscience – training the next generation of basic neuroscientists to embrace clinical research [108890/Z/15/Z]. A.S is supported by a Medical Research Council PhD Studentship in Precision Medicine with funding by the Medical Research Council Doctoral Training Programme and the University of Edinburgh College of Medicine and Veterinary Medicine. D.L.McC and R.E.M are supported by Alzheimer’s Research UK major project grant ARUK-PG2017B-10. J.M.W is supported by the Fondation Leducq.

