## Supplementary Methods for "An epigenetic predictor of death captures multi-modal measures of brain health"

**Methylation preparation in the Lothian Birth Cohort 1936**

Briefly, raw intensity data were background-corrected and normalised using internal controls. Manual inspection resulted in the removal of low quality samples presenting issues relating to bisulphite conversion, staining signal, inadequate hybridisation or nucleotide extension. Quality control analyses were performed to remove probes with low detection rate i.e. P < 0.01. Samples with a low call rate (samples with < 450,000 probes detected at p-values of less than 0.01) were eliminated. Samples were also removed if they had a poor match between genotype and SNP control probes, or incorrect DNAm-predicted sex.

For the current study, we applied a different quality control approach to the individuals that passed the screening process described above. This was to reduce missing CpG values, as recommended in Horvath’s epigenetic clock tutorial (https://dnamage.genetics.ucla.edu/). Raw DNAm IDAT files were read into R using the *minfi* package [1]. Data were normalised using the noob (normal-exponential convolution using out-of-band probes) method implemented by the preprocessNoob() function in *minfi* [1]. This method incorporates a background subtraction method and dye-bias normalisation. Briefly, the background subtraction method estimates background noise from out-of-band probes and remove it for each distinct sample, whereas the dye-bias normalisation uses a subset of control probes to estimate the dye bias. Noob-normalised methylation beta values were obtained using the getBeta() function in *minfi*.

**Protein measurement in the Lothian Birth Cohort 1936**

Plasma samples were analysed using a 92-plex proximity extension assay (Olink® Bioscience, Uppsala Sweden). In brief, 1 µL of sample was incubated in the presence of proximity antibody pairs linked to DNA reporter molecules. Upon binding of an antibody pair to their corresponding antigen, the respective DNA tails form an amplicon by proximity extension, which can be quantified by high-throughput real-time PCR. The data were pre-processed by Olink® using NPX Manager software.

**Brain imaging in the Lothian Birth Cohort 1936**

All imaging data (T1-, T2-, T2*-, and fluid attenuated recovery- and diffusion weighted scans) were acquired on a 1.5T GE (General Electric, Milwaukee, WI, USA) Signa HDx clinical scanner. Regional cortical volume was estimated using FreeSurfer v5.1. Following visual quality control to remove instances of aberrant surface estimation and gross segmentation errors, the *qcache* command was used to register each participant to the fsaverage surface, providing comparable volumetric measures at each of 327,684 vertices across the cortical mantle for all participants. After pre-processing, tract-averaged FA and MD were estimated in each of 12 white matter tracts using probabilistic neighbourhood tractography implemented in the TractoR package, using the BEDPOSTX/ProbTrackX algorithm, with two fibres modelled per voxel. The tracts were genu and splenium of the corpus callosum, bilateral anterior thalamic radiations, cingulum, uncinate, arcuate and inferior longitudinal fasciculi [2-5].

5.	Clayden, J.D., et al., *TractoR: magnetic resonance imaging and tractography with R.*
