## Supplementary Note 1 for "An epigenetic predictor of death captures multi-modal measures of brain health"

*Supplementary Note 1. Protocols for data collection of phenotypes in Hillary and Stevenson et al.*

1) Blood

Blood samples were obtained to assess:

- **Albumin** (g/L)
- **C-reactive protein** (mg/L)
- **Ferritin** (µg/L)
- **High-density lipoprotein cholesterol** (mmol/L)
- **Interleukin-6** (ng/ml)
- **Iron** (µg/L)
- **Low-density lipoprotein cholesterol** (mmol/L)
- **Total cholesterol** (mmol/L)
- **Triglycerides** (mmol/L)

2) Physical
   - **Body mass index** (kg/m^2^)
   - **Forced expiratory volume in 1 second (FEV1), forced vital capacity (FVC), forced expiratory ratio (FER (FEV1/FVC)), peak expiratory flow (PEF)** were all measured using a Micro Medical Spirometer, each the best of three.
   - **Grip strength** in both right and left hands (kg; measured using a North Coast Hydraulic Hand Dynamometer, JAMAR)

- **Height** (cm)
- **Weight** (kg)

3) Cognitive

- Six subsets of the Wechsler Adult Intelligence Scale (1) were included:
- **Digit Symbol Coding** was used to evaluate the speed of information processing. Participants enter a symbol according to a number-symbol code and complete as many as possible within a two-minute time-frame.
- **Symbol Search** also assesses speed of information processing. Participants examine a row of symbols to determine whether this row contains a pair of target symbols, and complete as many items as possible within an allotted time-frame.
- **Matrix Reasoning** assesses non-verbal reasoning. Participants are asked to examine patterns displayed in a matrix and identify the missing item based on this rule.
- **Letter-Number Sequencing** assesses working memory. Participants listen to increasingly long strings of numbers and letters and repeat these in exact numerical and alphabetical order.
- **Block Design** assesses constructional ability. In this test, participants reconstruct specific designs from blocks with a maximum of two minutes allotted per design task.
- **Backward Digit Span** assesses working memory. Participants are required to listen to increasingly long strings of numbers and repeat them backwards.
- Tests of prior (or crystallised) cognitive ability included the **Wechsler Test of Adult Reading** (2) and the **National Adult Reading Test (NART)** (3). These tests comprise 50 written words designed to test the participant’s vocabulary. The words have irregular spellings which do not possess regular pronunciation rules.
- **Mini-mental state examination (MMSE)** is a 30-point clinical questionnaire designed to measure cognitive impairment and screen for dementia (4). In this test, individuals are asked to answer questions relating to orientation of time, orientation to place, registration, attention, calculation, recall, language, repetition and complex commands. Scores ≤ 12 points indicate severe cognitive impairment, scores between 13-19 points are indicative of moderate impairment, scores between 20-24 points indicate mild cognitive impairment whereas scores ≥ 25 points corresponding to cognitive normal individuals.
- Three items from the Wechsler Memory Scale-III (5) were included:
- **Logical memory** I and II (total score) – These tests assess immediate and delayed verbal declarative memory respectively. Logical memory I entails the immediate recollection a given story which has 25 elements. The story is read aloud to the participant. Two stores are read with recall after each. The second story is read twice and participants are informed that they will have to recall the story later. Logical memory II involves recalling as much information as possible from the two stories which are read aloud in logical memory I.
- **Spatial span** is a test of non-verbal, spatial learning and memory. The participant watches a tester touching the top of a number of blocks in a spatial array with the aim of touching the same blocks in the same order. The task is repeated with the aim of touching the blocks in the reverse order.
- **Verbal paired associates** (total score) tests verbal learning and memory. Testers are asked to read a list of paired words in which some pairs have no obvious connection. They are then read the first of each pair and asked to recall the other. In total, there are 8 word pairs and the task is repeated in different orders four times. Following a delay and without the pairs of words being read again the task is repeated.
- **Simple and four choice reaction time (mean)** was used to assess speed of simple information processing. The tasks were administered using a stand-alone shallow rectangular box constructed for the UK Health and Lifestyle Survey (6). There are five response keys numbered 1,2,3,0 and 4. In the simple test, there are 8 practice tests followed by 20 test trials. The participant rests the second finger of the preferred hand on the 0 key. After a 0 appears on the screen, the participant presses the key as fast as possible. The mean of the 20 test trials are calculated. The four-choice version has 8 practice trials and 40 test trials. The participant rests the second and third fingers of the left and right hands on, respectively, the keys marked 1, 2, 3 and 4. After a number appears on the screen the participant presses the appropriate key as quickly as possible. Separate means are computed for correct and incorrect trials.
- **Inspection Time** was used is a two-alternative, forced choice, backward masking, visual discrimination task. It assesses the speed of elementary visual processing. The

inspection time task was replicated as closely as possible from a previous study (7), but with a longer instruction period and more practice trials. The participants were required to indicate which of two parallel, vertical lines of markedly different lengths was longer. The inspection time test was constructed, run, and analysed using E-Prime (Psychology Software Tools, Pittsburgh, PA). The stimulus lines were 5cm for the longer line and 2.5cm for the shorter line. The lines were joined at the top with a 2.5cm crossbar. Ten trials were carried out at each of 15 durations (rounded to the nearest millisecond): 6, 12, 19, 25, 31, 37, 44, 50, 62, 75, 87, 100, 125, 150, and 200. Participants indicated the position of the longer line by pressing 1 or 2 on the number pad of a computer keyboard. The correctness of each response was noted.

- **Verbal fluency** assesses executive functioning. In this test, participants are asked to name as many words as possible beginning with the letters C, F and L. Participants are given one minute per letter. No proper names are allowed and any repeated words are counted once.
- **Age 11 IQ** was calculated from participants’ Moray House Test score, corrected for age in days at the time of testing, and then converted to an IQ score. The Moray House Test was the test taken by most of the LBC1936 participants aged 11 as part of the Scottish Mental Survey 1947. The participants re-sat the test at Wave 1 (age 70) using the same instructions and with the same time limit. The test has a variety of items including: following directions (14 items), same-opposites (11), word classification (10), analogies (8), practical items (6), reasoning (5), proverbs (4), arithmetic (4), spatial items (4), mixed sentences (3), cypher decoding (2), and other items (4).

4) Neuroimaging

- Measures of brain volume, grey matter, white matter and white matter hyperintensities were normalised to intracranial volume to control for potential confounding effects of head size.
- Diffusion-tensor imaging generated metrics of fractional anisotropy and mean diffusivity.
- Full details in Supplementary Methods.

5)	Proteomics

- Information pertaining to proteomic data in the LBC1936 are described in full in Supplementary Methods.

the cognitive state of patients for the clinician. Journal of psychiatric research.

1975;12(3):189-98.

5. Wechsler D. WMS-IIIUK administration and scoring manual. Psychological Corporation. 1998.
6. Deary IJ, Der G, Ford G. Reaction times and intelligence differences: A population-based cohort study. Intelligence. 2001;29(5):389-99.
7. Deary IJ, Simonotto E, Meyer M, Marshall A, Marshall I, Goddard N, et al. The functional anatomy of inspection time: an event-related fMRI study. NeuroImage. 2004;22(4):1466-79.
