## Supplementary Note 2 for "An epigenetic predictor of death captures multi-modal measures of brain health"

*Supplementary Note 2. The association of age-adjusted DNAm Pack Years and DNAm blood proteins with physical, cognitive, neuroimaging and protein traits in the LBC1936 study.*

**DNAm Pack Years**

*Physical*

In relation to physical and blood traits, age-adjusted DNAm Pack Years was positively associated with levels of interleukin-6 (β = 0.25, SE = 0.04, P = 1.6 x $10^{-9}$), C-reactive protein (β = 0.18, SE = 0.04, P = 4.3 x $10^{-5}$), creatinine (β = 0.13, SE = 0.04, P = 2.0 x $10^{-3}$) and triglycerides (β = 0.11, SE = 0.04, P = 0.02). As with DNAm GrimAge, only the association with triglyceride concentration failed to remain significant after adjusting for age 11 IQ (% attenuation: 42.08%).

Age-adjusted DNAm Pack Years was negatively associated with all four measures of lung function (β = [-0.11 to -0.23] SE = [0.03 to 0.04], P = [1.0 x $10^{-3}$ to 2.0 x $10^{-13}$]), levels of iron (β = -0.16, SE = 0.04, P = 1.0 x $10^{-3}$), low-density lipoprotein cholesterol (β = -0.10, SE = 0.04, P = 0.03), and height (β = -0.07, SE = 0.03, P = 0.04). Only the association with height failed to remain significant following controlling for childhood cognitive ability (% attenuation 34.33%).

*Cognitive*

Following adjustment for age 11 IQ, age-adjusted DNAm Pack Years was significantly associated with the g factor (β = -0.08, SE = 0.03, P = 0.03), lower scores on the digit symbol coding (β = -0.11, SE = 0.04, P = 0.01) and symbol search tasks (β = -0.10, SE = 0.04, P = 0.02) and an increased mean four choice reaction time (β = 0.12, SE = 0.04, P = 6.0 x $10^{-3}$).

*Neuroimaging*

Age-adjusted DNAm Pack Years was negatively associated with ratios of white matter volume:ICV (β = -0.25, SE = 0.04, P = 3.0 x $10^{-7}$), brain volume:ICV (β = -0.19, SE = 0.04, P = 6.0 x $10^{-5}$), grey matter volume:ICV (β = -0.16, SE = 0.04, P = 1.0 x $10^{-3}$). Furthermore, age-adjusted DNAm Pack Years was positively associated with the ratio of white matter hyperintensities:ICV (β = 0.16, SE = 0.04, P = 2.0 x $10^{-3}$). All associations remained significant after adjusting for age 11 IQ.

*Protein*

DNAm Pack Years was significantly associated with the levels of 30/92 Olink® proteins, 24 of these associations remained significant following adjustment for age 11 IQ (Supplementary File 5a).

**DNAm Adrenomedullin**

*Physical*

In relation to physical and blood traits, age-adjusted DNAm Adrenomedullin (ADM) was positively associated with levels of interleukin-6 ($\beta$ = 0.30, SE = 0.04, P = $8.6 \times 10^{-10}$), C-reactive protein ($\beta$ = 0.15, SE = 0.04, P = 0.01), higher body mass index ($\beta$ = 0.21, SE = 0.04, P = $1.7 \times 10^{-4}$) and weight ($\beta$ = 0.13, SE = 0.04, P = 0.01). All associations remained significant after adjusting for age 11 IQ.

Age-adjusted DNAm ADM was negatively associated with three measures of lung function: peak expiratory flow ($\beta$ = -0.12, SE = 0.04, P = 0.01), forced expiratory volume ($\beta$ = -0.10, SE = 0.03, P = 0.02), forced vital capacity ($\beta$ = -0.10, SE = 0.03, P = 0.02) and levels of iron ($\beta$ = -0.16, SE = 0.05, P = 0.01). All associations remained significant after adjusting for age 11 IQ.

*Cognitive*

Following adjustment for age 11 IQ, age-adjusted ADM was significantly associated with the g factor ($\beta$ = -0.10, SE = 0.04, P = 0.03).

*Neuroimaging*

Age-adjusted DNAm ADM was not significantly associated with any neuroimaging phenotype.

*Protein*

DNAm ADM was significantly associated with the levels of 16/92 Olink® proteins, 15 of these associations remained significant following adjustment for age 11 IQ (Supplementary File 5b).

**DNAm Beta-2 microglobulin**

*Physical*

Age-adjusted DNAm Beta-2 microglobulin (B2M) was positively associated with levels of interleukin-6 (β = 0.16, SE = 0.04, P = $3.0 \times 10^{-3}$).

*Cognitive*

Age-adjusted DNAm B2M was not significantly associated with any cognitive phenotype.

*Neuroimaging*

Age-adjusted DNAm B2M was negatively associated with the ratios of grey matter volume:ICV (β = -0.16, SE = 0.04, P = $7.0 \times 10^{-3}$) and brain volume:ICV (β = -0.16, SE = 0.04, P = $7.0 \times 10^{-3}$).

*Protein*

DNAm B2M was significantly associated with the levels of 6/92 Olink® proteins, 5 of these associations remained significant following adjustment for age 11 IQ (SIGLEC1, MSR1, IL12, SCARB2 and Beta NGF) (Supplementary File 5c).

**DNAm Cystatin C**

*Physical*

Age-adjusted DNAm Cystatin C was positively associated with levels of interleukin-6 ($\beta$ = 0.14, SE = 0.04, P = 5.0 x $10^{-3}$) and body mass index ($\beta$ = 0.14, SE = 0.04, P = 5.0 x $10^{-3}$).

Age-adjusted DNAm Cystatin C was negatively associated with levels of low-density lipoprotein cholesterol ($\beta$ = -0.14, SE = 0.04, P = 5.0 x $10^{-3}$), total cholesterol ($\beta$ = -0.12, SE = 0.04, P = 0.01), iron ($\beta$ = -0.14, SE = 0.04, P = 0.01) as well as forced vital capacity ($\beta$ = -0.08, SE = 0.04, P = 0.04). All associations remained significant following controlling for age 11 IQ.

*Cognitive*

After adjusting for age 11 IQ, age-adjusted DNAm Cystatin C was negatively associated with the g factor ($\beta$ = -0.09, SE = 0.03, P = 0.04).

*Neuroimaging*

Age-adjusted DNAm Cystatin C was negatively associated with the ratios of grey matter volume:ICV ($\beta$ = -0.20, SE = 0.04, P = 8.6 x $10^{-5}$), brain volume:ICV ($\beta$ = -0.20, SE = 0.04, P = 2.5 x $10^{-4}$) and white matter volume:ICV ($\beta$ = -0.14, SE = 0.04, P = 0.02). All associations were significant after correcting for age 11 IQ.

*Protein*

DNAm Cystatin C was significantly associated with the levels of 3/92 Olink® proteins, all of which remained significant following adjustment for age 11 IQ (G CSF, SIGLEC9 and MSR1) (Supplementary File 5d).

**DNAm Growth Differentiation Factor 15**

*Physical*

Age-adjusted DNAm Growth Differentiation Factor 15 (GDF15) was not associated with any physical phenotype.

*Cognitive*

Age-adjusted DNAm GDF15 was not associated with any cognitive phenotype.

*Neuroimaging*

Age-adjusted DNAm GDF15 was not associated with any neuroimaging phenotype.

*Protein*

DNAm GDF15 was not significantly associated with the levels of any Olink® protein (Supplementary File 5e).

**DNAm Leptin**

*Physical*

Age-adjusted DNAm Leptin was positively associated with levels of interleukin-6 (β = 0.25, SE = 0.07, P = 0.01) and body mass index (β = 0.24, SE = 0.04, P =0.01). After adjusting for age 11 IQ, the association between age-adjusted DNAm Leptin and body mass index did not remain significant (β = -0.21, SE = 0.07, P = 0.05; % attenuation: 14.71%).

Age-adjusted DNAm Leptin was negatively associated with grip strength in the left hand (β = -0.13, SE = 0.02, P = 0.04).

*Cognitive*

Before adjusting for age 11 IQ, age-adjusted DNAm Leptin was negatively associated with the Wechsler test of adult reading (β = -0.20, SE = 0.07, P = 0.04). However, this association was attenuated following adjustment for age 11 IQ (% attenuation: 36.48%).

*Neuroimaging*

Age-adjusted DNAm Leptin was negatively associated with the ratios of brain volume:ICV (β = -0.30, SE = 0.07, P = $9.0 \times 10^{-3}$) and grey matter volume:ICV (β = -0.23, SE = 0.08, P = 0.04). After adjusting for age 11 IQ, the association between DNAm Leptin and grey matter volume:ICV did not remain significant (β = -0.21, SE = 0.08, P = 0.08; % attenuation: 9.21%)

*Protein*

DNAm Leptin was significantly associated with the levels of 4/92 Olink® proteins, all of which remained significant following adjustment for age 11 IQ (Beta NGF, SCARB2, EFNA4 and MSR1) (Supplementary File 5f).

**DNAm Plasminogen activator inhibitor-1**

*Physical*

Age-adjusted DNAm Plasminogen activator inhibitor-1 (PAI1) was positively associated with levels of triglycerides (β = 0.28, SE = 0.04, P = 1.3 x $10^{-9}$), body mass index (β = 0.28, SE = 0.04, P = 1.5 x $10^{-9}$), levels of interleukin-6 (β = 0.27, SE = 0.04, P = 2.1 x $10^{-9}$), weight (β = 0.21, SE = 0.04, P = 8.2 x $10^{-7}$) and levels of C-reactive protein (β = 0.19, SE = 0.04, P = 6.4 x $10^{-5}$).

Age-adjusted DNAm PAI1 was negatively associated with three measures of lung function: forced viral capacity volume (β = -0.19, SE = 0.03, P = 7.6 x $10^{-9}$), forced expiratory volume (β = -0.18, SE = 0.03, P = 1.4 x $10^{-7}$), peak expiratory flow (β = -0.11, SE = 0.03, P = 0.01) as well as levels of low-density lipoprotein cholesterol (β = -0.21, SE = 0.04, P = 3.3 x $10^{-6}$), total cholesterol (β = -0.14, SE = 0.04, P = 3.0 x $10^{-3}$), high-density lipoprotein cholesterol (β = -0.12, SE = 0.04, P = 0.02) and grip strength in the left hand (β = -0.07, SE = 0.03, P = 0.03). Only the association between age-adjusted DNAm PAI1 and high-density lipoprotein cholesterol did not remain significant after adjusting for age 11 IQ.

*Cognitive*

After adjusting for age 11 IQ, age-adjusted DNAm PAI1 was not associated with any cognitive phenotype.

*Neuroimaging*

Age-adjusted DNAm PAI1 was negatively associated with the ratios of brain volume:ICV (β = -0.24, SE = 0.04, P = 8.2 x $10^{-7}$), grey matter volume:ICV (β = -0.21, SE = 0.05, P = 6.4 x $10^{-5}$) and white matter volume:ICV (β = -0.16, SE = 0.04, P = 3.0 x $10^{-3}$). After adjusting for age 11 IQ, all associations remained significant.

*Protein*

DNAm PAI1 was significantly associated with the levels of 21/92 Olink® proteins, 19 of these associations remained significant following adjustment for age 11 IQ. Additionally, a further three proteins were significantly associated with DNAm PAI1 after adjusting for age 11 IQ (G CSF, KYNU and THY1) (Supplementary File 5g).

**DNAm Tissue inhibitor of metalloproteinases 1**

*Physical*

Age-adjusted DNAm Tissue inhibitor of metalloproteinases 1 (TIMP1) was positively associated with levels of C-reactive protein (β = 0.15, SE = 0.04, P = 3.0 x $10^{-3}$), interleukin-6 (β = 0.16, SE = 0.04, P = 3.0 x $10^{-3}$), creatinine (β = 0.11, SE = 0.04, P = 0.02) and body mass index (β = 0.10, SE = 0.04, P = 0.04).

Age-adjusted DNAm TIMP1 was negatively associated with levels of iron (β = -0.17, SE = 0.04, P = 3.0 x $10^{-3}$), forced expiratory volume (β = -0.10, SE = 0.03, P = 9.0 x $10^{-3}$), peak expiratory flow (β = -0.10, SE = 0.03, P = 0.01). All associations remained significant after adjusting for age 11 IQ.

*Cognitive*

Age-adjusted DNAm TIMP1 was not associated with any cognitive phenotype.

*Neuroimaging*

Age-adjusted DNAm TIMP1 was not associated with any neuroimaging phenotype.

*Protein*

DNAm TIMP1 was significantly associated with the levels of 16/92 Olink® proteins, 15 of these associations remained significant following adjustment for age 11 IQ. Additionally, a further two proteins were significantly associated with DNAm TIMP1 after adjusting for age 11 IQ (EFNA4 and LAYN) (Supplementary File 5h).
