## Supplementary Figures 1-7 for "An epigenetic predictor of death captures multi-modal measures of brain health"

***Supplementary Figure 1***

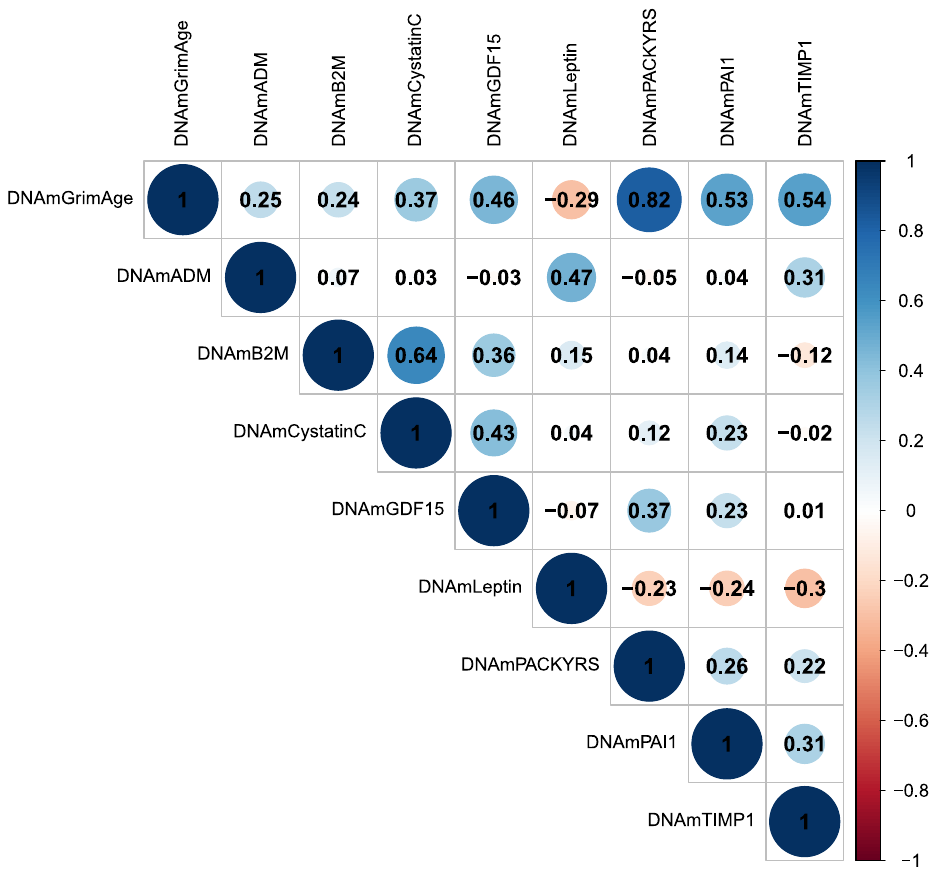

***Supplementary Figure 2***

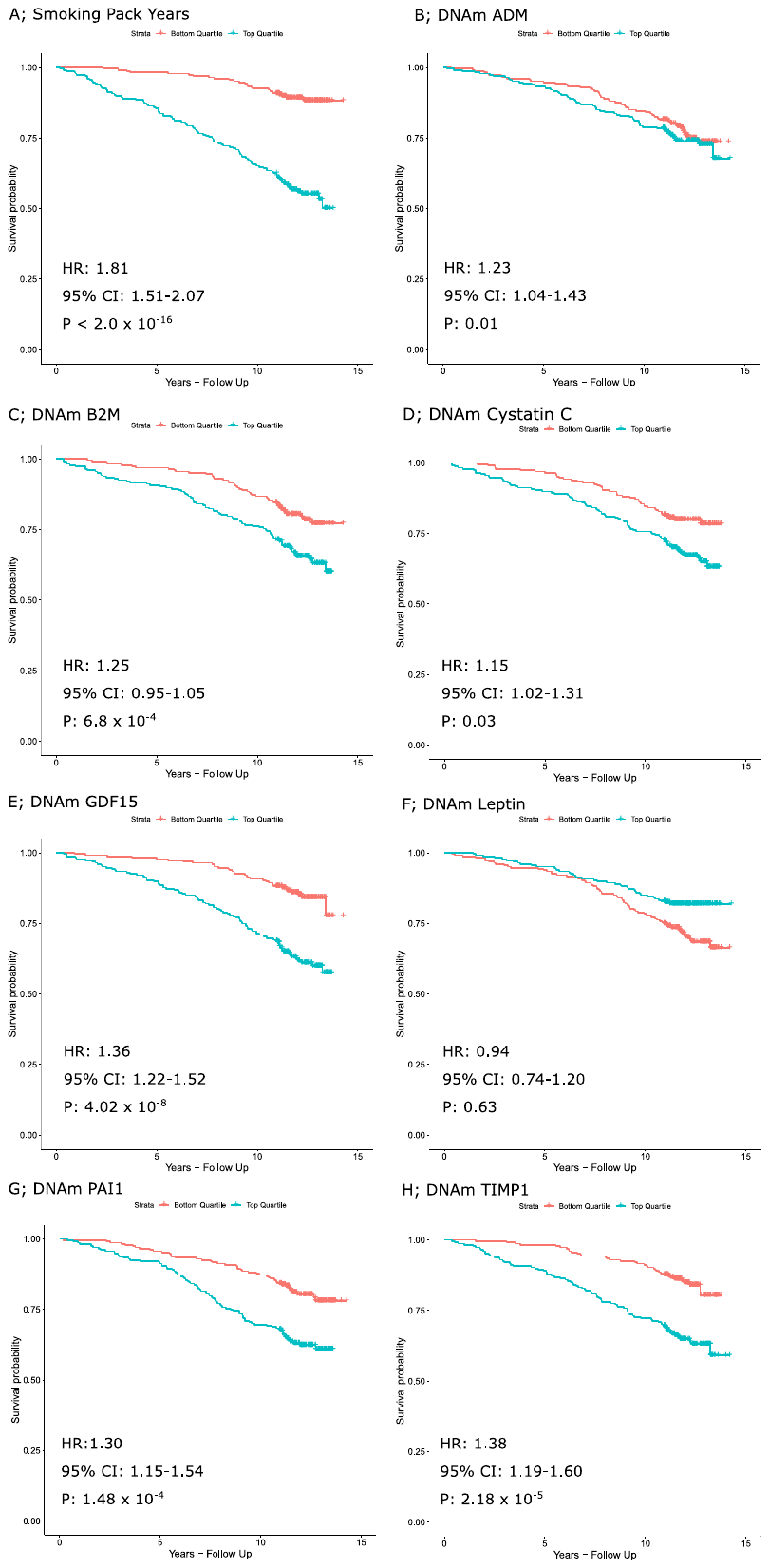

***Supplementary Figure 3***

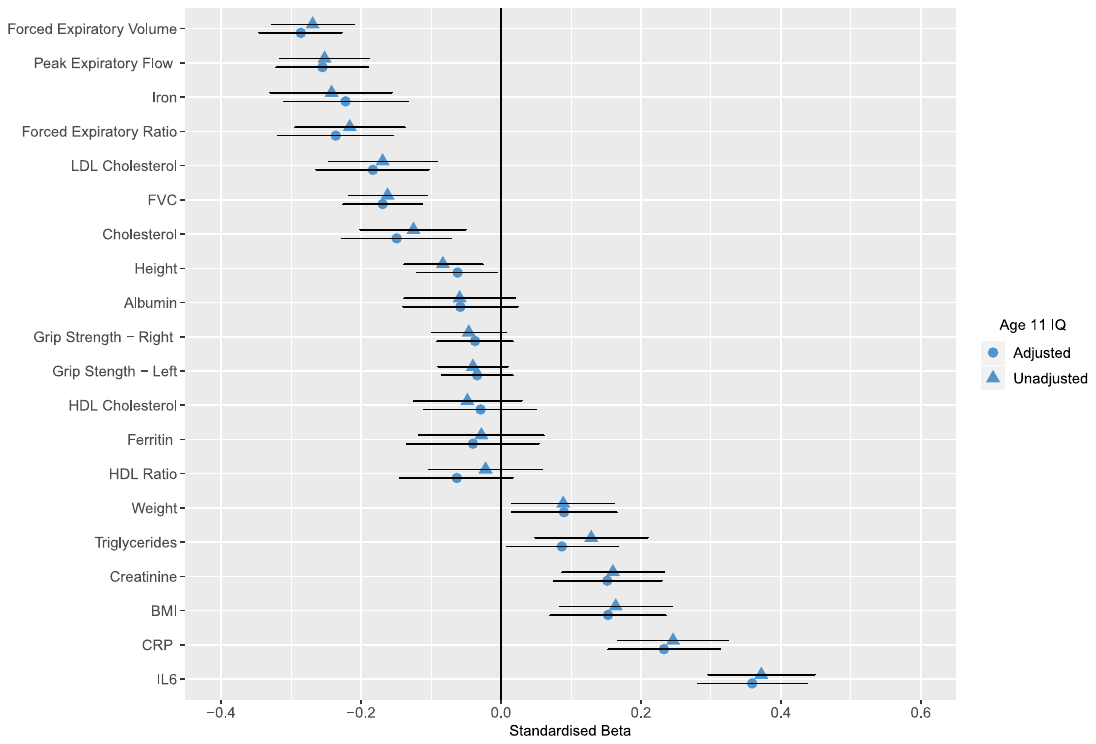

***Supplementary Figure 4***

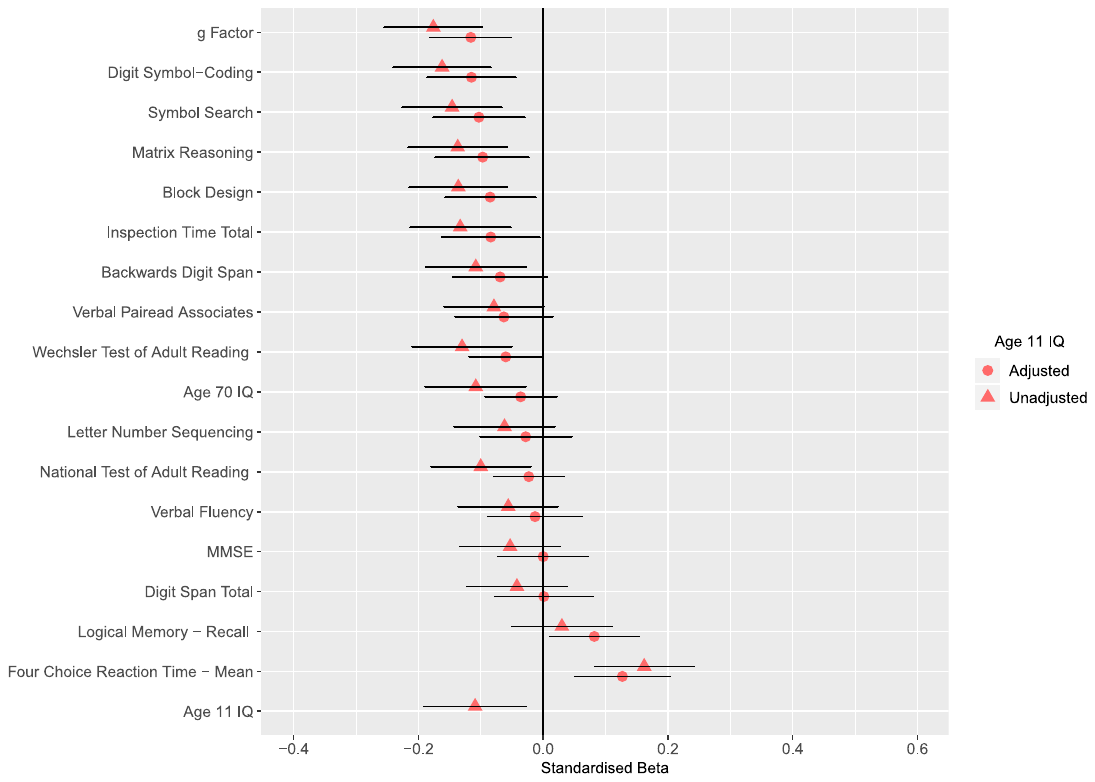

***Supplementary Figure 5***

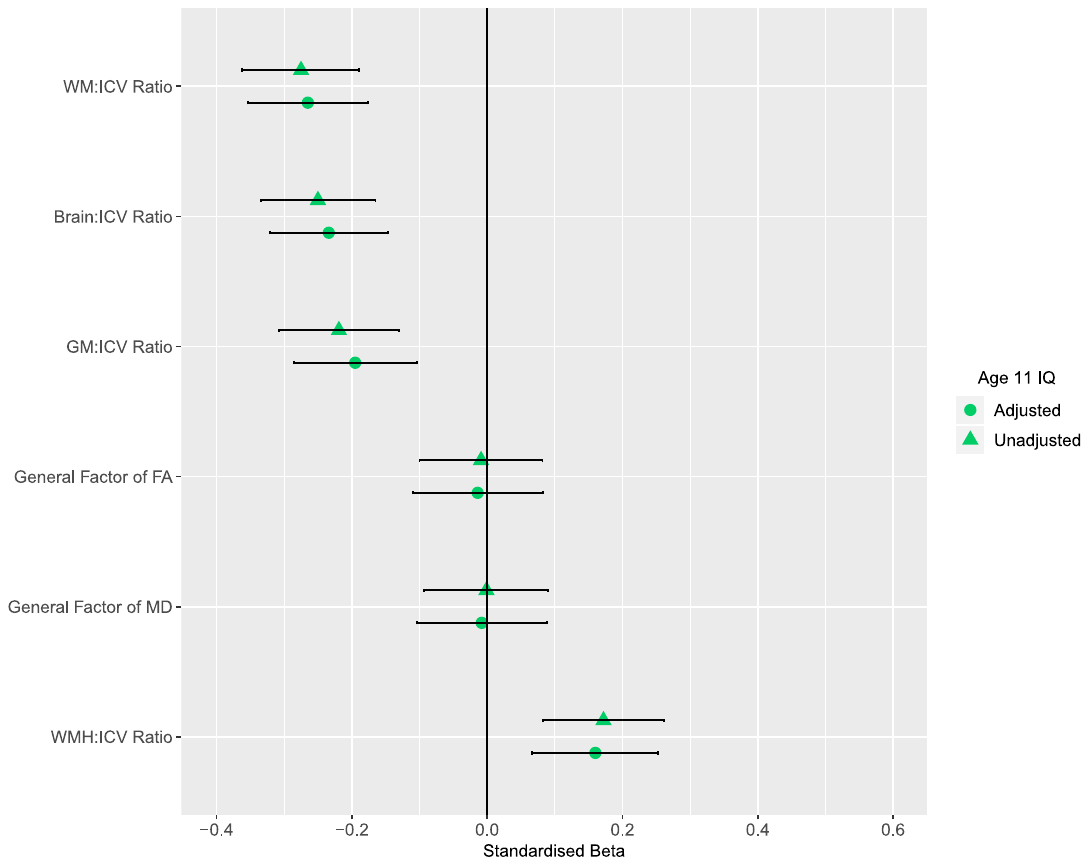

***Supplementary Figure 6***

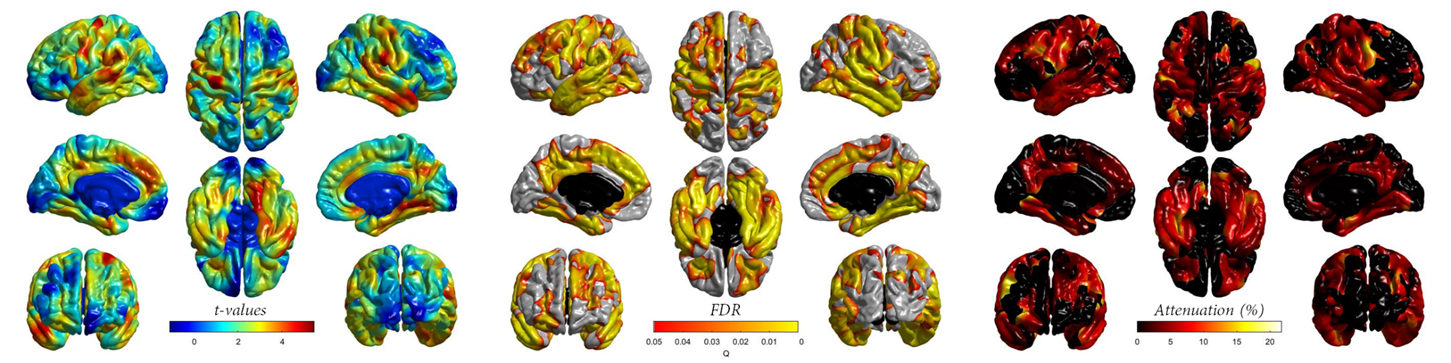

**Supplementary Figure 6. Cross-sectional association between age-adjusted DNAm GrimAge and regional cortical volume in the LBC1936, corrected for age 11 IQ.** *Left panel:* t-values indicate the magnitude of the negative association (values have been flipped for visualisation purposes). *Centre Panel*: Corresponding FDR-corrected P values indicate the spatial distribution of significant associations. *Right Panel:* Degree of attenuation of t-values at initially FDR-significant loci before and after correcting cortical volumetric-DNAm GrimAge associations for age 11 IQ. FDR (false discovery rate).

***Supplementary Figure 7***

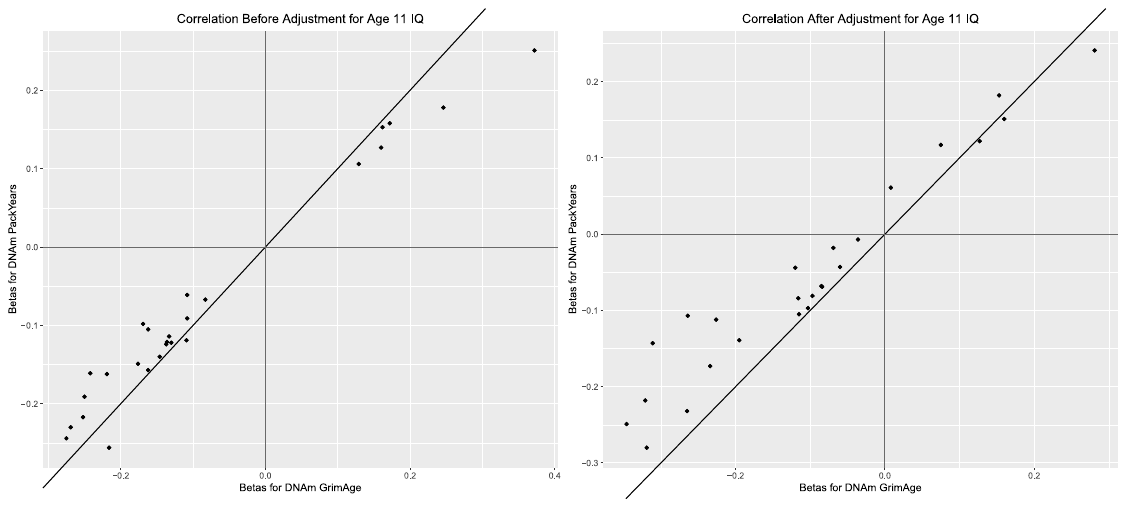
