## Supplementary File 1 for "An epigenetic predictor of death captures multi-modal measures of brain health"

| *Supplementary File 1. Summary of demographics and phenotypic variables in LBC1936 at Wave 2.* BMI (body mass index), HDL (high-density lipoprotein), ICV (intracranial volume), IQ (intelligence quotient), LDL (low-density lipoprotein). |
| --- |

| ***Trait*** | ***Count (%)*** | | ***n*** |
| --- | --- | --- | --- |
| Sex (% Female) | 379/796 (47.61%) | | 709 |
| ApoE4 carrier (% carrier) | 226/754 (29.97%) | | 682 |
|  | ***Mean*** | ***SD*** | ***N*** |
| Age (years) – wave 1 | 69.60 | 0.80 | 906 |
| DNAm GrimAge (years) – wave 1 | 67.40 | 5.20 | 906 |
| Age (years) – wave 2 | 72.54 | 0.70 | 709 |
| DNAm GrimAge (years) – wave 2 | 69.98 | 4.88 | 709 |
| *Cognitive* |  |  |  |
| Age 11 IQ (age 11 Moray House test score, corrected for age in days, then converted to IQ) | 100.69 | 15.37 | 666 |
| Age 70 IQ | 101.1 | 14.01 | 703 |
| Block Design | 33.68 | 10.09 | 707 |
| Four Choice Reaction Time – Mean | 0.65 | 0.09 | 708 |
| Backwards Digit Symbol | 7.80 | 2.30 | 709 |
| Digit Symbol | 56.27 | 12.29 | 706 |
| Inspection Time Total | 111.11 | 11.79 | 689 |
| Logical Memory Recall | 37.06 | 9.02 | 709 |
| Letter Number Sequencing | 10.91 | 3.08 | 709 |
| Matrix Reasoning | 13.19 | 4.97 | 707 |
| Mini Mental State Examination | 28.72 | 1.44 | 708 |
| National Adult Reading Test | 34.4 | 8.23 | 707 |
| Spatial Span Total | 14.68 | 2.75 | 706 |
| Simple Reaction Time – Mean | 0.27 | 0.05 | 708 |
| Symbol Search | 24.6 | 6.12 | 707 |
| Verbal Fluency | 43.15 | 13.05 | 708 |
| Verbal Paired Associates | 6.36 | 2.05 | 691 |
| Wechsler Test of Adult Reading | 40.96 | 7.02 | 707 |
| g factor | 0.01 | 0.99 | 703 |
| *Neuroimaging* |  |  |  |
| Brain volume:ICV ratio | 0.68 | 0.02 | 540 |
| Grey Matter volume:ICV ratio | 0.33 | 0.02 | 540 |
| White Matter volume:ICV ratio | 0.33 | 0.02 | 540 |
| White Matter Hyperintensities volume:ICV ratio | 0.009 | 0.01 | 540 |
| General Factor of Mean Diffusivity | 5.00E-17 | 1 | 550 |
| General Factor of Fractional Anisotropy | 1.60E-17 | 1 | 550 |
| *Physical* |  |  |  |
| Albumin 35-50 g/L | 43.73 | 2.9 | 704 |
| BMI (kg/m^2^) | 27.9 | 4.38 | 709 |
| Creatinine 60-120 µmol/L | 80.34 | 24.55 | 706 |
| C-reactive Protein (mg/L) | 2.92 | 5.34 | 704 |
| Ferritin 20-300 µg/L | 122.27 | 136.12 | 564 |
| Grip Strength - Right Hand | 28.64 | 9.39 | 707 |
| Grip Strength - Left Hand | 27.56 | 9.44 | 707 |
| Height (cm) | 166.52 | 8.95 | 709 |
| HDL Ratio (Total: HDL Cholesterol) | 3.76 | 1.12 | 705 |
| HDL Cholesterol 0.9 - 1.4 mmol/L | 1.45 | 0.42 | 705 |
| Interleukin-6 (ng/ml) | 130.83 | 85.87 | 707 |
| Iron 10-32 µmol/L | 17.07 | 5.67 | 561 |
| LDL Cholesterol 2-5 mmol/L | 2.93 | 1.06 | 703 |
| Weight (kg) | 77.53 | 14.31 | 709 |
| Total Cholesterol (mmol/L) | 5.14 | 1.17 | 705 |
| Triglycerides 0.8 - 2.1 mmol/L | 1.67 | 0.83 | 705 |
| Forced Expiratory Volume in 1 second (FEV1) | 2.3 | 0.69 | 702 |
| Forced Vital Capacity (FVC) | 3.11 | 0.9 | 702 |
| Forced Expiratory Ratio (FEV1/FVC) (FER) | 77.09 | 11.82 | 702 |
| Peak Expiratory Flow (PEF) | 346.49 | 128.26 | 702 |
